# Prior movement of one arm facilitates motor adaptation in the other

**DOI:** 10.1101/2022.11.22.517483

**Authors:** M. Gippert, S. Leupold, T. Heed, I. S. Howard, A. Villringer, V. V. Nikulin, B. Sehm

## Abstract

Many movements in daily life are embedded in motion sequences that involve more than one limb, demanding the motor system to monitor and control different body parts in quick succession. During such movements, systematic changes in the environment or the body might require motor adaptation of specific segments. However, previous motor adaptation research has focused primarily on motion sequences produced by a single limb, or on simultaneous movements of several limbs. For example, adaptation to opposing force fields is possible in unimanual reaching tasks when the direction of a prior or subsequent movement is predictive of force field direction. It is unclear, however, whether multi-limb sequences can support motor adaptation processes in a similar way. In the present study, we investigated whether reaches can be adapted to different force fields in a bimanual motor sequence when the information about the perturbation is associated with the prior movement direction of the other arm. In addition, we examined whether prior perceptual (visual or proprioceptive) feedback of the opposite arm contributes to force field-specific motor adaptation. Our key finding is that only active participation in the bimanual sequential task supports pronounced adaptation. This result suggests that active segments in bimanual motion sequences are linked across limbs. If there is a consistent association between movement kinematics of the linked and goal movement, the learning process of the goal movement can be facilitated. More generally, if motion sequences are repeated often, prior segments can evoke specific adjustments of subsequent movements.

**Significance statement:** Movements in a limb’s motion sequence can be adjusted based on linked movements. A prerequisite is that kinematics of the linked movements correctly predict which adjustments are needed. We show that use of kinematic information to improve performance is even possible when a prior linked movement is performed with a different limb. For example, a skilled juggler might have learned how to correctly adjust his catching movement of the left hand when the right hand performed a throwing action in a specific way. Linkage is possibly a key mechanism of the human motor system for learning complex bimanual skills. Our study emphasizes that learning of specific movements should not be studied in isolation but within their motor sequence context.

## 1 Introduction

Many movements in daily life are embedded in motion sequences that involve more than one limb. Interaction between two arms in frequently repeated sequences is usually fast and effortless. For example, a juggler is able to transfer juggling balls rapidly and precisely between two hands. Due to variability in single motor segments, the juggler has to be able to adjust movements accordingly. In the current study, we investigate how the motor system is able to learn such intricate mechanisms to adjust movements in bimanual sequences. We show that once a motor sequence is learned, kinematic information from the beginning of the sequence can be used to modify later segments.

If two or more movements in a sequence are repeated in the same order many times, the individual motor elements seem to be linked together in a single motor action (Diedrichsen and Kornysheva, 2015; Verwey et al., 2015). Thus, a motor element which is strongly linked in a sequence can influence prior and following motor segments (Hansen et al., 2018). If a reach is linked to a prior movement of the same arm, kinematic characteristics of that prior movement can even facilitate motor adaptation (Howard et al., 2012). In other words, information from a preceding same-limb movement can be used to adjust the following movement accordingly.

Motor adaptation studies involving a single force field have shown that an internal model of the motor dynamics to counteract external forces is acquired over time (Anwar et al., 2011). When multiple force fields are experienced, interference problems arise. Simple visual cues that indicate perturbation direction are ineffective in eliciting motor adaptation to opposing force fields (e.g., Cothros et al., 2009). In contrast, cues that allow overcoming the interference of multiple force fields are related to the motor plan (Hirashima and Nozaki, 2012; Howard et al., 2017; Sarwary et al., 2015; Sheahan et al., 2016; Wainscott et al., 2005) or the sensory state of the arm (Howard et al., 2013; Green and Labelle, 2015; Sarwary et al., 2013; Crevecoeur et al., 2022). Such cues are thought to enable cue-specific motor memory formation and retrieval by putting the sensorimotor system in the right preparatory state (Howard et al., 2020). Linked movements in particular seem to be effective cues because the representation of the entire motor sequence is specific to each perturbation direction and thus allows the creation of separate sensorimotor memories for each force field. In addition, sensory same-arm movement cues – passive or visual prior movements – are as effective for field-specific adaptation as actively performed linked movements (Howard et al., 2012). This suggests that perceptual information that implies the sensory consequences of same-arm movement execution can be linked to the active target reach.

Bimanual motor adaptation research, however, has been focused primarily on simultaneously executed movements rather than sequential movements (Tcheang et al., 2007; Howard et al., 2008; Kadota et al., 2014; Nozaki et al., 2006). Therefore, despite its relevance for human motor behavior, it is yet unknown if a prior opposite-arm movement can serve as an effective cue for force field specific adaptation. Successful adaptation would indicate that distinct sequential segments of bimanual sequences can be linked together. During juggling this would mean that, for instance, specific kinematics of the throwing action of one arm could be linked to specific adjustments during the catching motion of the other arm.

In the present study we, thus, investigated linkage of movements of two arms. In addition to the replication of previous unimanual findings, we examined whether, and to what extent, a prior movement of the opposite arm could facilitate adaptation of the following movement in a force field interference task. To answer what aspects are key for establishing a link between such movements, we tested whether prior sensory information from the other arm (vision or proprioception) could be used as an effective cue for force field specific adaptation.

## 2 Materials and Methods

Our study design, including sample sizes, hypotheses and main analysis plan, was pre-registered on OSF (https://osf.io/qy9rn). Data and analysis scripts we used to arrive at the results we present here will be made available upon publication. In total, 68 right-handed volunteers aged 18-35 years (34 female, 34 male) participated in our study. We excluded 4 participants from analysis (see below), thus our sample comprised 64 (32 female, 32 male, *M*_*age*_ = 26, *SD*_*age*_ = 4.3) participants. All participants had normal or corrected-to-normal vision and were free of any known neurological, perceptual and motor impairments and disorders. The experiment was approved by the local ethics committee of the University of Leipzig. Participants gave informed written consent prior to the experiment.

All participants were required to make reaching movements in a Kinarm Exoskeleton Lab (see Figure 1; for a video of the task see OSF repository). The Kinarm Exoskeleton Lab is a robotic device that can measure movements of the arms in the horizontal plane at a sampling rate of 1000 Hz. The robot can apply forces to the arms and provide visual feedback in a two-dimensional augmented virtual environment.

**Figure 1:**
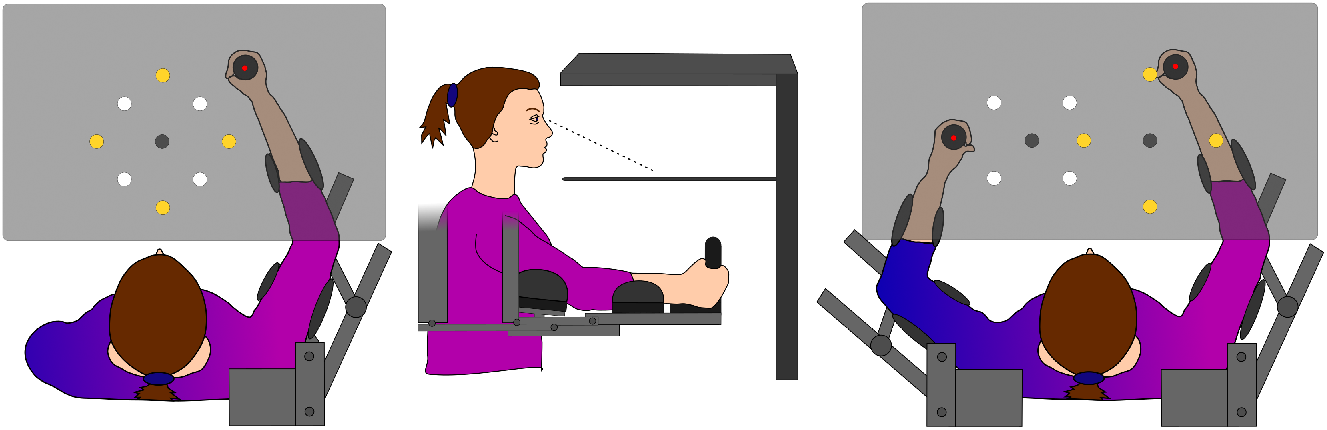
Experimental setup. Kinarm Exoskeleton Robot Lab. All possible target positions are shown. The screen is displayed transparent here; however, in the experiment participants were not able to see their arms. Left: Setup unimanual groups. Middle: Side view. Right: Setup bimanual groups.

### 2.1 Experimental Design

Participants were randomly assigned to one of five groups (see Figure 2). Each group performed reaches to targets in the Kinarm. In two of the five groups, participants were required to move only their right arm (unimanual groups), while the other three groups incorporated both hands in subsequent reaching movements (bimanual groups). In the unimanual groups, three targets were displayed during each trial: the cue, middle and final target (see Figure 3A). The middle target was individually calibrated to be at the position of the hand when the elbow was flexed 90°and the shoulder angle was 60°. In the bimanual groups, there were four targets: the cue, middle-left, middle-right and final target. The distance between the two middle targets was 18 cm, and the midway point between them was fixed at 90° elbow and 60° shoulder angle for each participant. Each trial consisted of one or two active reaches. The final reach in each trial was the same for all experimental groups: participants made a reaching movement from the middle or middle-right target to the final target with their right hand.

**Figure 2:**
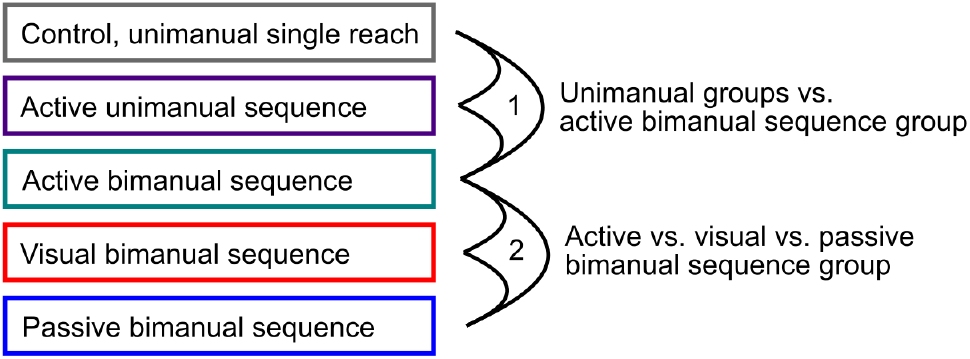
Experimental groups and comparisons. All groups and planned comparisons between groups to answer 1) whether and to what extent the effectiveness of a prior arm movement to allow force field specific adaptation generalizes to bimanual sequences and 2) whether prior sensory information from the other arm in one modality (vision or proprioception) can be used instead of active movement for force field specific adaptation.

**Figure 3:**
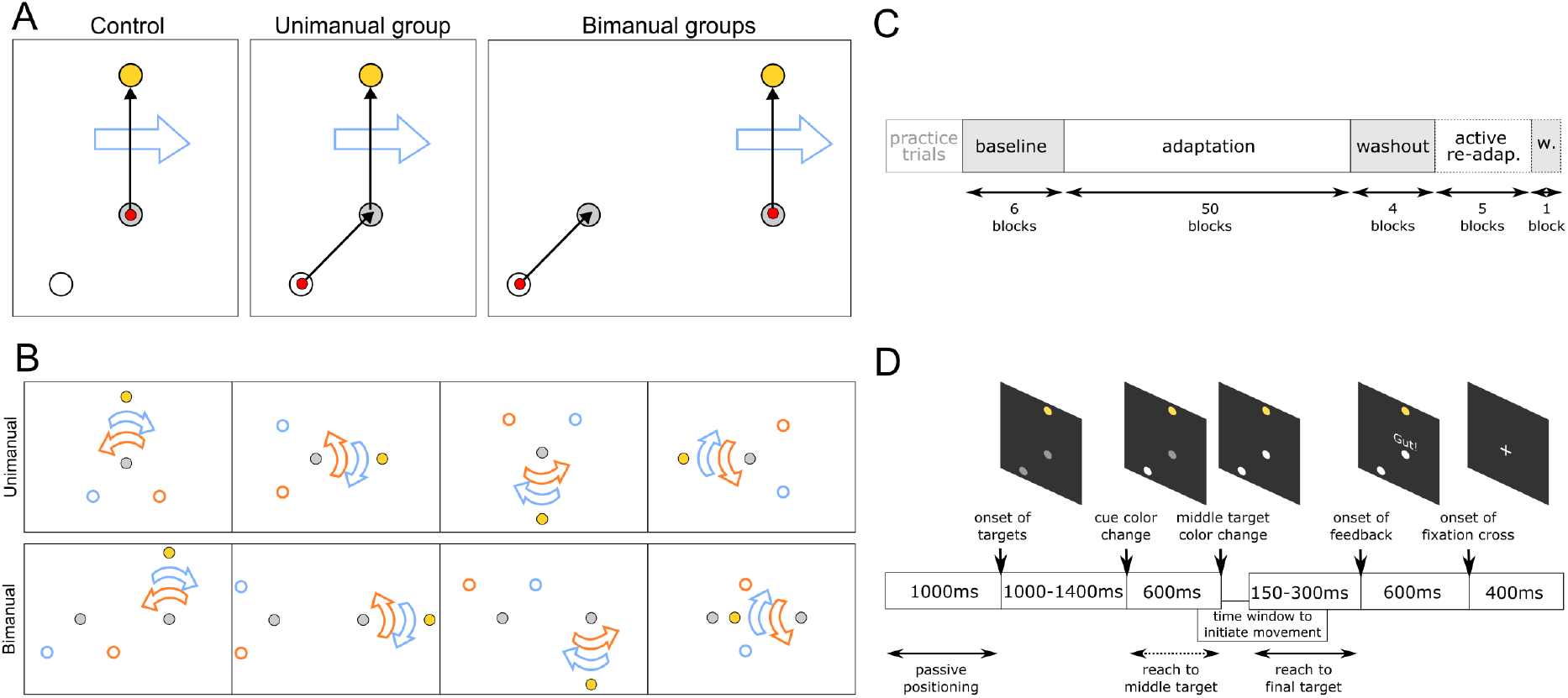
Trial Design and experimental schedule. A) Exemplary trial. Yellow dot - final target; grey dot - middle target(s); white dot - cue; red dot - hand position at the beginning of a trial; black arrow - desired reaching path during the trial; blue arrow - force field direction. In the bimanual groups the left hand reach was either performed actively, passively or only visually displayed. B) All cue final target combinations with arrows representing force field direction. For half of the participants the relationship between cue position and force field direction was the other way around. Yellow dot - final target; grey dot(s) - middle target(s); white dot with blue/orange border - cues; blue/orange arrow - force field direction. C) Experimental flow. re-adap. = re-adaptation. w. = washout. Phases with dashed lines are only executed in bimanual groups. D) Trial sequence. Reach to middle target was not (actively) performed in all groups.

In all groups, there were four possible final target positions: 12 cm to the right, left, up or down from the middle (-right) target. There were two possible cue positions for each final target position (see Figure 3B). The distance between cue and middle (-left) target was 10 cm. The cursor displaying current hand position was red and 0.5 cm in diameter. All targets were 1.25 cm in diameter. The final target was yellow. All other targets were initially grey and changed their color to white during the trial (see below).

All groups went through three experimental phases: baseline, adaptation, and washout (see Figure 3C). Bimanual groups went through two additional experimental phases thereafter, namely readaptation and another washout phase. These latter two phases aimed at assessing potential transfer of learning from different bimanual sequence group conditions to a bimanual group condition which involved actively moving both arms.

During trials in the adaptation and re-adaptation phases, a velocity-dependent curl field was present between the middle (-right) and the final targets. This force field systematically perturbed the right hand’s movements. The force field started with a ramp up time of 100 ms once the right hand was more than 2 cm away from the midpoint of the middle target and stopped once the final target had been reached. The cue’s location in relation to the final target was uniquely associated with the direction of the force field. Half of the participants learned the association between a positive angle between cue and final target and a clockwise (CW) force field, the other half between a positive angle and a counterclockwise (CCW) force field to control for any kinematic or biomechanical advantages of a specific combination. The association between the sign of the angle and the direction of the force field was fixed for each participant and did not change during the experiment.

The forces experienced during adaptation trials were perpendicular to movement direction and proportional to reaching velocity:

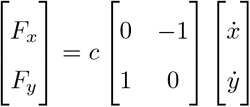

where the constant c was set to -13 Ns/m or +13 Ns/m depending on the location of the cue (Howard 2012). The resulting force field was CW or CCW, respectively. There was never a force field between the cue and the middle target(s) and there was never any force field present in baseline or washout trials.

In 11% of trials, chosen randomly throughout the trial sequence, the Kinarm forced movements to be straight by means of a force channel, that is, a force field that resembles a straight channel with impenetrable walls on its sides. In these trials, no curl force field was present. We term these trials clamp trials, as they restrict participants movements to a straight trajectory while they attempt to counteract the expected (but absent) force field perturbation. The Kinarm measures the compensatory forces applied by the participant against the channel walls during the clamp trial, allowing us to quantify any possible feed-forward learning, which would be expressed in the compensation of the expected (but absent) force field. Due to technical constraints, clamp trials did not always generate a strong enough force channel. In this case, the hand broke through the virtual wall and deviated from the straight trajectory. We excluded these trials, and so analysis of the last four adaptation block was based on 422 of 512 trials.

At the beginning of the experiment, participants were briefly familiarized with the task. They then performed 6 baseline, 50 adaptation and 4 washout blocks. The bimanual groups performed 6 additional blocks – 5 re-adaptation and one additional washout block (see below). Each block consisted of 16 normal and 2 clamp trials. In total, participants performed at least 1080 trials. There were short breaks approximately every 200 trials and a 5 min break at the halfway point. Number and size of targets, timing, angles and force field strength were derived from the literature (e.g., Howard et al., 2012). At the end of the experiment, participants filled out pencil-paper questionnaires that asked them about potential strategies used in the experiment. In addition, we asked whether they had recognized a specific pattern between force field direction and cue position.

#### Group 1: control, unimanual single reach

We excluded and replaced two participants in the control group, because they had detected the relationship between force field direction and position of targets and reported to have used this knowledge explicitly to move faster through the force field. The sample used for analysis was 20 (10 female, 10 male, *M*_*age*_ = 26.55, *SD*_*age*_ = 5.17). We recorded EEG in the unimanual groups for another study, which is why the sample size was larger for unimanual than bimanual groups.

At the beginning of each trial, a white fixation cross on black background was displayed and the right arm of participants was moved to the middle target by the robot (see Figure 3D). This passive positioning took 1000 ms. Then, all targets (cue, middle & final) and the hand position cursor were displayed. After a random time drawn from a uniform distribution ranging from 1000 ms to 1400 ms, the cue changed color from grey to white. This event was meaningful in the other experimental groups; in contrast, control group participants were not instructed to do anything yet. After 600 ms the middle target changed color from grey to white, which was the go-signal for participants to reach to the final target. Once it was reached, feedback about the movement speed was displayed right above the middle target. If the movement time from 2 cm away from the middle to the final target was in between 150 and 300 ms, feedback (in German) was ’good’; if it was outside this range ’too fast’ or ’too slow’, respectively. The feedback was displayed for 600 ms. Finally, a white fixation cross was shown for 400 ms before the next passive positioning started for the next trial. When half the trials of a block were completed the inter-trial-interval was 4 s instead of 400 ms.

Trials were immediately aborted and repeated within the same block when the cursor was not in the middle target when the color changes of the targets took place. If participants left the middle target earlier than 100 ms before or later than 500 ms after the go-signal, the trial was marked unsuccessful and repeated at a random position within the current block. Timing, feedback and repetition criteria, as introduced here, were the same for all groups.

For two participants in the control group the time between cue color change and middle target color change was set to 400 ms instead of 600 ms. Performance of these two was similar to other participants in the group and so we included them in our analysis.

#### Group 2: active unimanual sequence

There were 20 participants in the unimanual sequence group (10 female, 10 male, *M*_*age*_ = 25.9, *SD*_*age*_ = 4.45). Unlike in the control group the hand of the participants was moved to the position of the cue and not the middle target during the passive positioning. The color change of the cue was the indicator for participants to move from the cue to the middle target. Participants were instructed to try to reach the middle target approximately when it changed color and subsequently reach from the middle to the final target. The aim was to perform two separate straight reaches but to pause in the middle target as short as possible. Feedback at the trial’s end referred only to the movement time from middle to final target. Trials were aborted when the cursor was not in the cue when the first color change occurred and when the cursor left the middle target too early or too late (see control group).

Four participants had to perform faster cue-middle target reaches because the time between cue color change and target color change was only 400ms. Their performance was similar to that of other participants and so we included them in the data analysis.

#### Group 3: active bimanual sequence

The active bimanual sequence group comprised 8 participants (4 female, 4 male, *M*_*age*_ = 27.25, *SD*_*age*_ = 4.23). Due to large effect sizes observed in prior research (Howard et al., 2012; Sheahan et al., 2016), we chose this sample size for all two hands groups. Although there was one more visual target in the two hand groups, timing of color changes was exactly the same as in the one-hand groups. The two middle targets changed color at the same time, 600ms after the cue color change.

At the beginning of a trial, the Kinarm robot moved both the right and the left arm; the left cursor to the cue position and the right cursor to the middle-right target. The color change of the cue was the signal for the participants to move the left arm to the middle-left target. Like in the unimanual sequence group, they were instructed to reach the middle-left target approximately when the middle targets changed color. The goal was to finish the left hand movement and subsequently reach with the right hand to the final target. To keep the trial abortion and repetition criteria consistent with the unimanual groups, participants had to have both cursors in their respective starting positions once the color change of the cue indicated the start of the left arm reach. Trials were also repeated when the right hand left the middle-right target more than 100ms too early or more than 500ms too late. Like in the unimanual sequence group, force fields and force channels (for clamp trials) were only ever present for the second reach, but never for the left hand. Feedback displayed after the movements was only about the right hand movement speed from the middle-right to the final position.

There were 6 additional blocks in the bimanual groups. After the last washout block, participants performed 5 blocks in an active bimanual sequence re-adaptation condition and subsequently one final active washout block. These blocks were identical to those of the first adaptation and washout phases.

#### Group 4: passive bimanual sequence

Our sample comprised 8 participants (4 female, 4 male, *M*_*age*_ = 25.25, *SD*_*age*_ = 2.92). We excluded and replaced two participants because they employed explicit strategies, which they reported in our debriefing questionnaire.

Like in the active bimanual sequence group, the left arm was positioned at the cue and the right arm at the middle-right target at the beginning of a trial. However, participants did not see a cursor at the position of the left hand and they did not actively move their left arm during the three main phases of the experiment. Instead, participants were instructed to keep their left arm relaxed while the Kinarm moved the left hand from cue to middle-left target following a minimum jerk trajectory after the cue color change. This passive reach of the left arm started 100ms after the go-signal and took 550 ms to mirror an average active reach and the preceding reaction time. Participants were instructed to start their right hand reach to the final target once they felt that the passive movement of the left hand finished. They were told that the color change of the middle targets did not always occur at the same time in relation to the passive left arm movement to discourage participants to discount the passive arm movement and only pay attention to the color change as a start signal. After the experiment, we asked participants whether they noticed that the end of the passive left arm movement always coincided with the middle targets color change and whether they had used this color change as a strategy to initiate their right arm reach.This was the case for one participants; we evaluate this point in the Discussion.

After the main three phases, participants performed five re-adaptation blocks and one washout block with the same instructions as the active bimanual sequence group. They experienced the same force field directions as in the adaptation phase; however, now they had to actively move the left hand. This post-test assessed whether there was any transfer of force field adaptation that may have taken place in the experiment’s prior phases from passive to active left hand movement.

#### Group 5: visual bimanual sequence

There were 8 participants in the visual bimanual sequence group (4 female, 4 male, *M*_*age*_ = 24, *SD*_*age*_= 2.73). During the main 3 phases of the experiment, participants did not move their left arm. At the start of each trial, only the right hand was positioned to the middle-right target. Once the targets were displayed, there was, however, also a red cursor at the cue position. After the cue color change, this cursor moved to the middle-left target with the same motion dynamics as the passive movement in the passive bimanual sequence group. Participants were instructed to start their right hand reach once the red cursor reached the middle-left target. They were also asked not to use the middle target color changes as a go-signal but instead focus on the moving red cursor. Three participants paid attention to the color changes; we address this point in the Discussion.

Like the other bimanual groups, participants performed 6 blocks of active bimanual movements at the end of the experiment to assess learning transfer.

### 2.2 Data analysis

We pre-processed data in Matlab (R2021a). The Kinarm measured angles of the elbow and shoulder joints. We low-pass filtered this data with a cutoff at 10 Hz and added hand velocity, acceleration and commanded forces.

#### Maximal perpendicular error (MPE)

We performed our main analysis in Python (3.7) using the libraries numpy (Harris et al., 2020), pandas (McKinney et al., 2010), scipy (Virtanen et al., 2020), scikit-learn (Pedregosa et al., 2011), as well as matplotlib (Hunter, 2007) and seaborn (Waskom, 2021) for plotting. We excluded all aborted and repeated trials. Our first outcome measure was the maximal perpendicular error (MPE), defined as the signed maximal deviation in cm from the straight line between middle (-right) and final target of the right arm trajectory and reflected adaptation performance. A positive value denoted that participants had exhibited a curved trajectory in the direction of the force field. The MPE was defined between 2 cm away from the midpoint of the middle target and the end of the final target. We excluded trials when it was obvious that participants had started the reach towards an incorrect target (13 across all samples).

#### Force field compensation (FFC)

Our second outcome measure was force field compensation (FFC) in clamp trials. To calculate FFC, we extracted force data within a 150 ms time window centered on the time of peak velocity. Next, we calculated the ideal force profile which would have counteracted the missing force field based on movement velocity. The measured force against the channel walls were linearly regressed on the ideal force profile with the intercept forced to zero. We defined FFC as the slope of the regression multiplied with 100%.

#### Statistical analysis

The focus of our study was on motor adaptation learning and differences in this aspect between groups. We quantified current motor adaptation performance with the MPE in normal trials and FFC in clamp trials. We averaged MPE and FFC over block and calculated the mean and standard error over participants for each group (see Figures 5A&B, 6A&B).

To measure the degree of adaptation each participant exhibited, we calculated three dependent variables. First, we subtracted the average MPE of the first adaptation block from the average of the last two adaptation blocks for each participant (MPE change adaptation). A negative MPE change adaptation value indicates that straighter reaches occurred at the end of the adaptation phase compared to the beginning. The more negative the value, the greater was the performance improvement. Second, we calculated the difference of the average MPE of the last baseline and first washout block (MPE change baseline/washout). A negative value means that a participant made systematically more curved reaches counteracting the experienced force fields in the washout block than in the baseline block. Consequently, the more negative the value, the bigger was the force field after-effect. This reaching behavior indicates that participants adapted to the force field and exhibited after-effects when the force field was removed (e.g., Gandalfo, 1996). Third, we calculated the average FFC for each participant in the last four blocks in the adaptation phase (FFC final adaptation). A FFC final adaptation value of 100% would indicate that participants perfectly adjusted their reaches to the force fields.

To answer our research questions, we performed planned comparisons between groups (see Figure 2). To assess differences in motor adaptation due to a same or opposite arm prior movement, we compared all three dependent variables between the control, uni-, and bimanual active sequence groups (see Figure 3). For each group comparison we calculated the difference of the mean of the dependent variable. Next, we permuted group labels and computed the resulting means’ difference. We repeated this 5000000 times or until the number of exact possible permutations was reached. The *p*-value was defined as the proportion of sampled permutations where the absolute difference was greater than the absolute observed difference. Following the same rationale, we compared all dependent variables within-group against zero (indicating no performance change) to assess whether group performance improved or worsened over time. For each family of permutation tests, we adjusted the *p*-value using the Bonferroni-Holm correction. We defined a family of tests as tests evaluating the same dependent measure (1 family = 11 tests).

To investigate whether sensory information in one modality of the opposite arm during the prior movement is sufficient for adaptation to occur, we compared performance measures between the three bimanual groups and within the groups. In addition, based on the observation that the learning curves of the active uni- and bimanual sequence groups differed, we investigated the slope of adaptation in both groups at the beginning of the adaptation phase (first 10 blocks) to identify potential learning differences at an early stage of the experiment. We performed a linear mixed effects analysis in R using afex (Singmann et al., 2016). We entered block number and group membership as well as the interaction term as fixed effects. As random effects, we included by-participant random slopes and intercepts. We employed the Kenward-Roger method to obtain *p*-values.

We used an explorative approach to examine performance in the re-adaptation phase in the bimanual sequence groups. We were interested to see if any transfer of learning occurred from a passive/visual to an active bimanual sequence. Due to the small number of blocks in the re-adaptation phase, we did not look at performance improvement within this phase but rather compared performance in this phase to performance at the end of adaptation. We averaged the MPE of the last five adaptation blocks of each participant and subtracted the average MPE of the re-adaptation phase (MPE re-/adaptation change). We again performed within and between permutation tests with this value.

To investigate to what extent the three measures, MPE change adaptation, MPE change baseline/washout and FFC final adaptation, reflect the same underlying factor, we calculated Pearson’s correlation between two measures each across participants.

## 3 Results

All participants performed reaches from a middle position to final targets. Opposing force fields perturbed reaches during the (re-)adaptation phase of the experiment. Figure 4 depicts all trajectories from middle (-right) to final target of relevant blocks of all participants. At the end of the adaptation phase, participants in the active sequence groups made straighter reaches between the middle and final targets than at the beginning of adaptation, when the force field had just been introduced. Moreover, when the force field was removed, strong after-effects were evident as curving of reach trajectories in the direction of the former force fields, due to participants being prepared to counter the force field they had previously encountered. Adaptation and after-effects indicate that participants adapted their reaches to the respective force fields.

**Figure 4:**
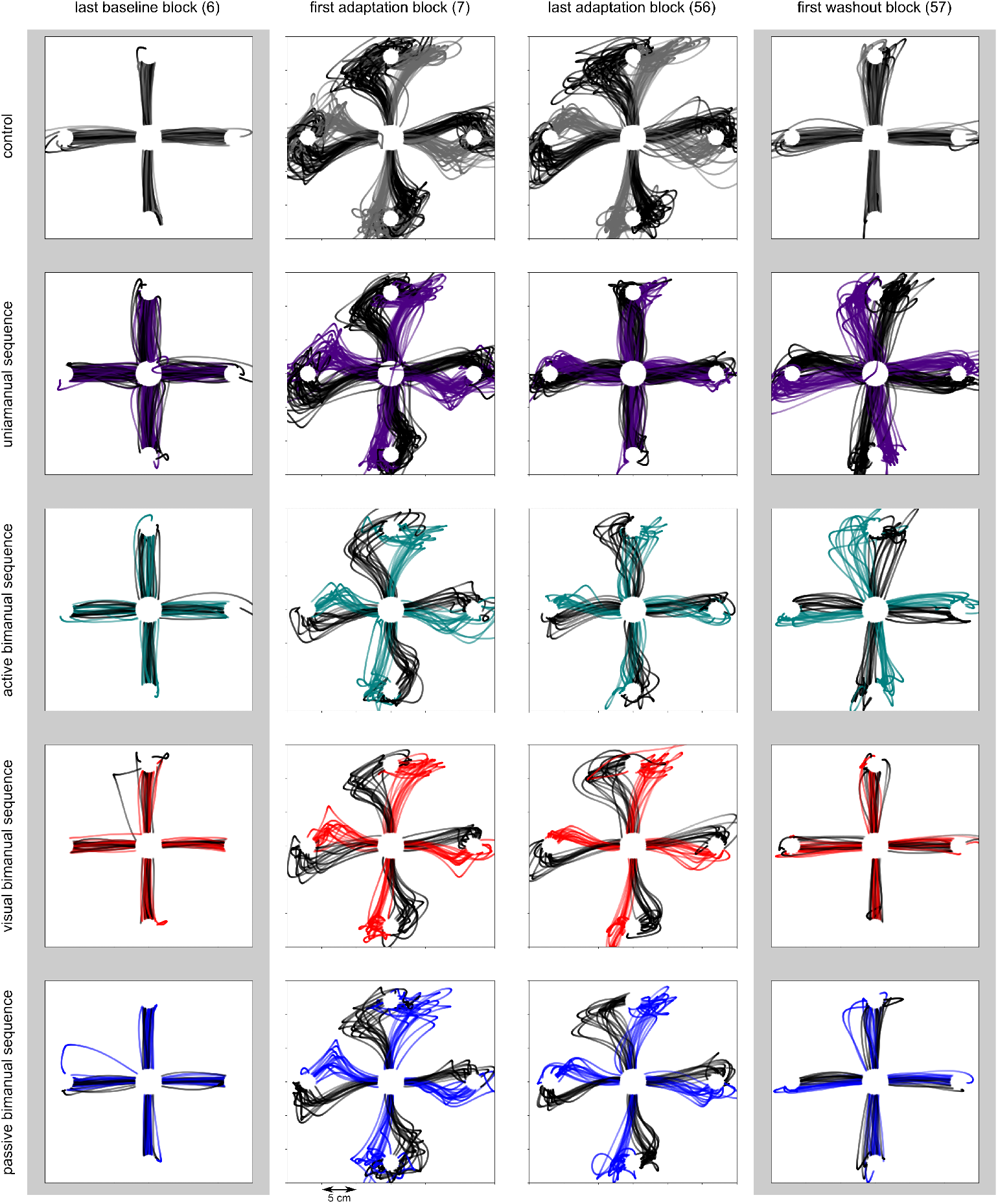
Reaching trajectories. Single trial trajectories of all participants in selected blocks of the experiment.

**Figure 5:**
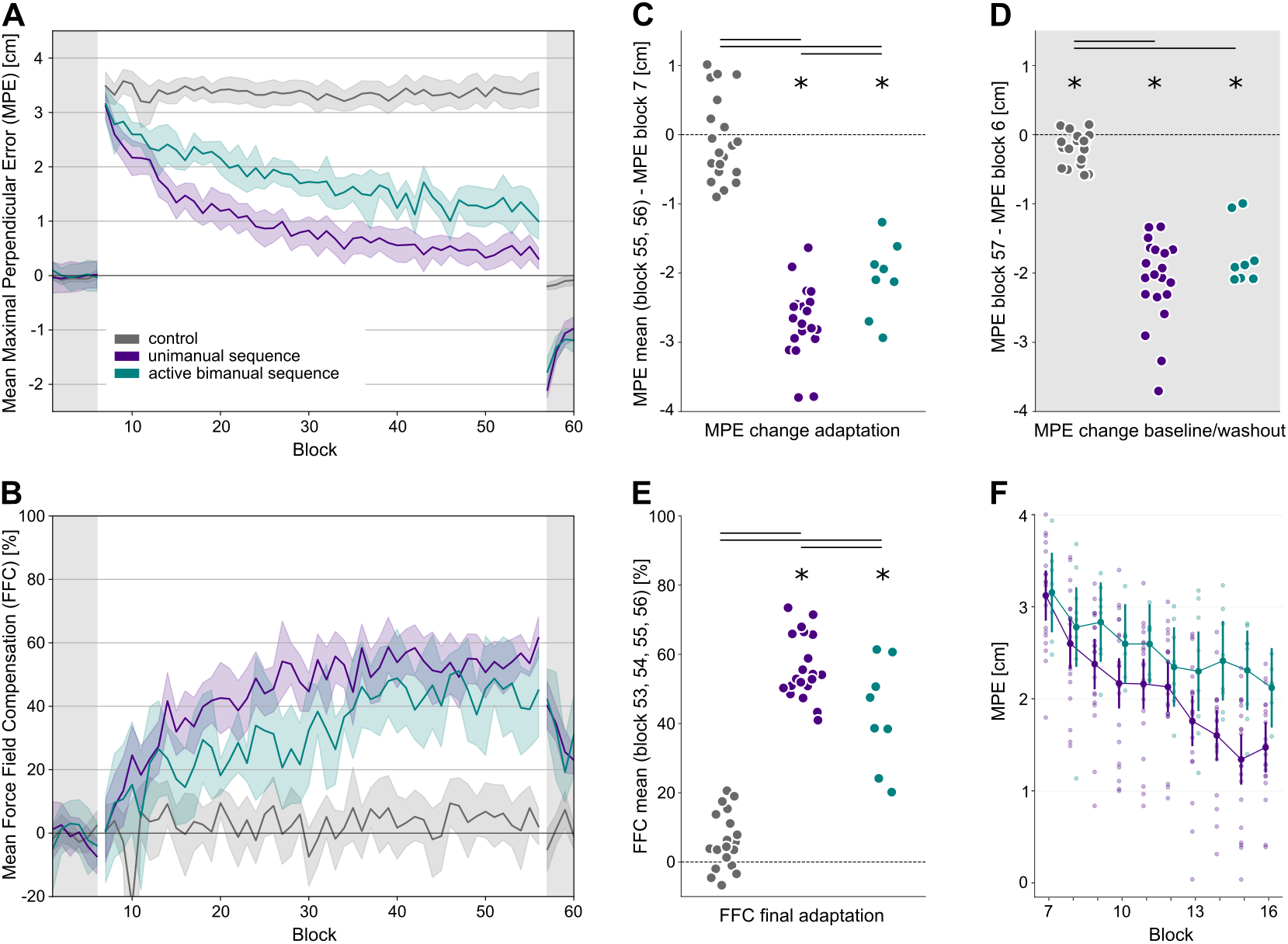
Unimanual groups vs. active bimanual sequence group. A) MPE averaged over participants and trials within one block. Force fields were present from block 7 to block 56 (white background). Error bands depict SEs across participants. B) FFC averaged over participants and trials within one block. C) Each dot depicts the difference between average performance in the first and last two adaptation blocks of one participant. Lines denote significant differences between groups *p <* .05. Stars mark significant within group effects *p <* .05. D) MPE differences between the first washout and the last baseline block. E) Average FFC in the last 4 adaptation blocks. F) MPE in the first 10 adaptation blocks; each small dot depicts average performance per block of one participant; larger dots display group averages; error bars depict 95% confidence intervals.

**Figure 6:**
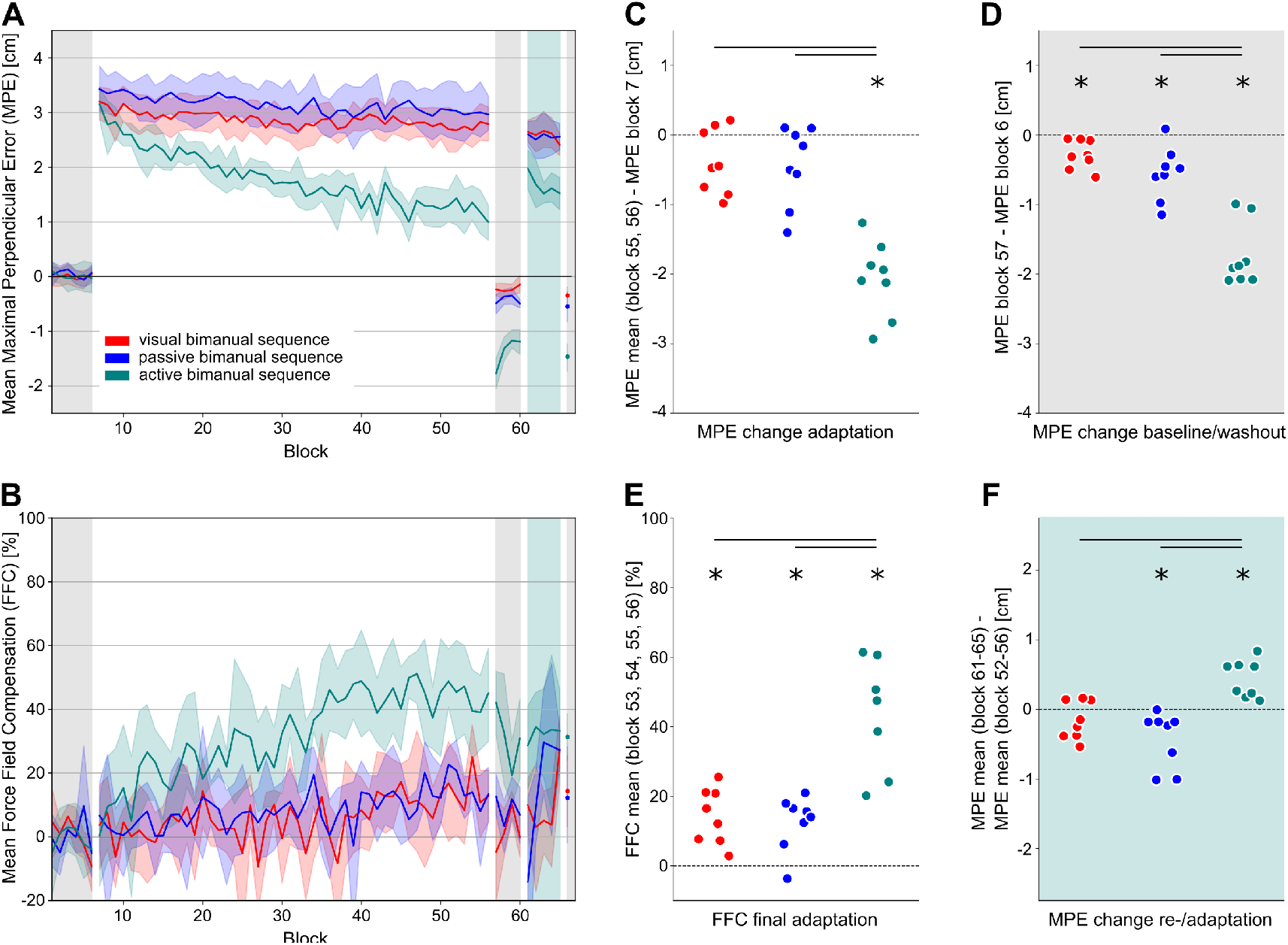
Active vs. visual vs. passive bimanual sequence group. A) MPE averaged over participants and trials within one block. Force fields were present from block 7 to block 56 (white background) and block 61 to block 65 (green background). All participants performed an active bimanual sequence in blocks 61 to 66. Error bands depict SEs across participants. B) FFC averaged over participants and trials within one block. C) Each dot depicts the difference between average performance in the first and last two adaptation blocks of one participant. Lines denote significant differences between groups p *<* .05. Stars mark significant within group effects p *<* .05. D) MPE differences between the first washout and the last baseline block. E) Average FFC in the last 4 adaptation blocks. F) MPE differences between the re-adaptation phase and the last 5 blocks of the adaptation phase.

### Unimanual groups vs. active bimanual sequence group

First, we asked whether a prior movement with the opposite arm can be used as an effective cue for learning force field specific adaptation. In addition, we investigated the degree of adaptation in comparison to same arm prior movements as a cue. To this end, we compared MPE changes and FFC final adaptation to zero (no performance change) within the control, unimanual and bimanual sequence group as well as across groups.

For each participant, we subtracted the average MPE of the first adaptation block from the average of the last two adaptation blocks to obtain a measure of the MPE change during the adaptation phase. MPE change during the adaptation phase was not significant in the control group (*p* = 0.5887; see Figure 5C). Control participants did not improve in counteracting the forces over the course of the adaptation phase. In contrast, performance of both the unimanual (*p* = 1.7e-05) and the active bimanual sequence group (*p* = 0.0469) improved over the adaptation phase. Improvement was greater in both sequence groups compared to the control group (*p*_*unimanual*_ *<* 1e-07; *p*_*bimanual*_ = 3.2e-06) and greater in the unimanual compared to the bimanual sequence group (*p* = 0.043). These results imply that prior movements of the opposite arm can setup the sensorimotor system in a way that allows force field specific adaptation. After repeated exposure to interfering forces during reaching to a target, movement kinematics of the prior opposite arm movement seem to be represented together with specific motor actions which allow counteracting the forces. This linkage of two movements of two arms seems to be less strong than linkage between two movements of one arm.

In addition, we assessed MPE change from the baseline to the washout phase by subtracting the average MPE of the last baseline block from the first washout block. MPE baseline/washout changes were present in all groups (*p*_*control*_ = 0.003; *p*_*unimanual*_ = 1.7e-05 ; *p*_*bimanual*_ = 0.043; see Figure 5D). This means that all groups showed some bias to curve their reaches in the direction from where the force field was coming from before. The changes were greater in the two sequence groups compared to the control group (*p*_*unimanual*_ *<* 1e-07; *p*_*bimanual*_ = 3.2e-06). There was no significant difference between the two sequence groups (*p* = 0.132). These results demonstrate that movements of two arms can be linked.

To confirm our findings with a measure of feedforward adaptation, we calculated FFC final adaptation by averaging forces measured in clamp trials in the last four blocks in the adaptation phase. FFC comparisons revealed the same pattern as MPE changes in the adaptation phase: FFC final adaptation was different from zero in both sequence groups but not in the control group (*p*_*unimanual*_ = 1.7e-05; *p*_*bimanual*_ = 0.043; *p*_*control*_ = 0.080; see Figure 5E). Stronger adaptation was observed in both sequence groups compared to the control group(*p*_*unimanual*_ = 0; *p*_*bimanual*_ = 6.4e-06). Finally, the unimanual sequence group compensated more over the course of the adaptation phase than the bimanual sequence group (*p* = 0.032). In sum, the result patterns are consistent with the notion that prior movement kinematics of the opposite arm can indeed serve as effective cues for force field specific motor adaptation. Yet, two of three dependent measures indicate that final adaptation is stronger in the uni-compared to the bimanual group.

To investigate whether this difference in adaptation occurs during early learning in the adaptation phase we performed a mixed model with the between groups factor group (uni-vs. bimanual sequence) and within group factor block (block 7-16). We found main effects for group (*F*_*1,26*_ = 4.65, *p* = .040) and block (*F*_*9,234*_ = 37.87, *p <* .001) as well as an interaction between group and block (*F*_*9,234*_ = 4.39, *p <* .001; see Figure 5F). Participants in the unimanual sequence group had a steeper learning curve than participants in the bimanual sequence group during early learning. This result indicates that linking of movements over body parts might be slower than linking movements within one body part.

#### Active vs. visual vs. passive bimanual sequence group

To investigate whether the perception of specific opposite arm movements without active execution allows motor adaptation, we tested for effects of adaptation (MPE changes and FFC final adaptation) in each group separately as well as across groups. MPE changes in the adaptation phase were neither significant for the visual (*p* = 0.0625) nor the passive (*p* = 0.0703) bimanual sequence group (see Figure 6C). Moreover, the active group improved their performance to a greater extent than the visual (*p* = 0.0012) and the passive (*p* = 0.0022) group. Finally, the change in performance was not different between the visual and passive group (*p* = 0.8382). These results suggest that perception of movement in one sensory modality, in the absence of active movement, does not sufficiently allow force field specific adaptation.

However, MPE changes from the baseline to the washout phase were evident in all groups (*p*_*visual*_ = 0.0430; *p*_*passive*_ = 0.0469; see Figure 6D). This indicates that some learning does occur in the sensory groups even though it is not evident in the adaptation phase. The changes were greater in the active compared to the visual (*p* = 0.0012) and passive (*p* = 0.0044) groups, highlighting that sensory information of the opposite arm can not be used as a substitute for active movement in this paradigm. No difference was observed between the visual and the passive group (*p* = 0.1032).

The overall result pattern was equivalent for FFC final adaptation (see Figure 6E). All groups showed some compensation (*p*_*visual*_ = 0.0430; *p*_*passive*_ = 0.0469), but it was stronger in the active compared to the visual (*p* = 0.0076) and the passive (*p* = 0.0025) group. The visual and passive groups did not differ in their FFC final adaptation (*p* = 0.6706). In total, these results confirm that movement directions of visual or passive prior movement with the other arm are not easily linked with motor actions needed to counteract a specific force field. Information about visual or passive movement kinematics of another limb might not be readily utilized to adjust movement plans.

#### Transfer

Finally, we asked whether the hidden learning in the visual/passive conditions could be transferred to an actively performed two arm motor sequence. To this end we compared the MPE from the end of adaptation to the MPE during the re-adaptation phase. We observed performance changes in the passive (*p* = 0.0273) and active (*p* = 0.0273) bimanual sequence groups, but not in the visual group (*p* = 0.1563; see Figure 6F). Participants in the active bimanual sequence group performed worse during the re-adaptation phase than at the end of the adaptation phase. This result is likely caused by the washout phase in between the adaptation and re-adaptation phase, in which participants re-adapted to an environment without force fields. Participants in the passive group, however, were able to reduce their MPE in the re-adaptation blocks compared to the last adaptation blocks. It is an open question whether this result is due to a motor learning mechanism or due to the need of the participants to stabilize their arm movements more when both arms have to be actively controlled. Comparisons between groups revealed a difference in MPE re-/adaptation change between the active and visual group (*p* = 0.0039) as well as between the active and passive group (*p* = 0.0009). The MPE was not different between the visual and passive groups (*p* = 0.1414). Overall, the transfer results further support the conclusion that initial learning was limited in the visual and passive group, making it difficult for transfer of learning to occur.

#### Correlation between measures

In order to investigate whether dynamics were similar across our different measures, we calculated a correlation between them. All correlations between MPE change adaptation, MPE change baseline/washout and FFC final adaptation across participants were high and in the expected direction (see table 1), suggesting that all variables measured the same underlying construct.

**Table 1:** Correlations between measures. Pearson’s r correlations between dependent variables.

|  | MPE change adaptation | MPE change baseline/washout |
| --- | --- | --- |
| MPE change baseline/washout | 0.8381 |  |
| FFC final adaptation | -0.8485 | -0.7881 |

## 4 Discussion

Fine-tuning movements to reduce discrepancies between motor command and sensory feedback is a key mechanism of motor learning. In this study, we investigated whether linking sequential movements of two arms can support motor adaptation to opposing force fields. We report two main findings. First, we found that prior movements of the opposite-arm enabled adaptation to opposing force fields. This finding demonstrates that learning of a movement can be influenced by a linked movement of the other arm. Specifically, if there are consistent relations between kinematics of different arm movements in a motion sequence, the motor system can take advantage of this information to adjust movements accordingly. Second, visual and proprioceptive opposite-arm signals in the absence of an active movement were significantly less effective than active reaches, highlighting that actively using both arms is a key requirement in linking sequential, bimanual movements. In addition, we replicated previous findings showing that active same-arm prior movements facilitate adaptation, whereas stationary visual cues, though indicative of force field direction, do not (e.g., Howard et al., 2012).

### Unimanual vs bimanual sequence learning

Our findings indicate that formation of separate motor memories in a force field interference task is not only possible when distinct perturbations are encountered in unimanual movement sequences, but also in active, bimanual movement sequences. Notably, the bimanual movement chains were less effective than unimanual chains, evident in both, the smaller reduction of trajectory curvature as measured with the MPE in the adaptation phase and smaller forces applied against the expected perturbation as indicated by the FFC measure. Furthermore, comparison of the adaptation curves for unimanual and bimanual sequences during early learning indicated that adaptation in the bimanual context was not only weaker but also slower than in the unimanual context. Thus, chained movements of a single limb seem to be more readily linked and represented together than movements across limbs. Several factors may explain this difference.

First, if single movement elements of a sequence are difficult to perform, it is more likely that they are represented separately as discrete actions (Rand and Stelmach, 2000). This is also reflected by an increased response time when alternating between different hands in bimanual serial reaction time tasks (Bhakuni and Mutha, 2015; Trapp et al., 2012). Using different limbs within one sequence adds complexity, requiring more coordination and attention to execute the movement sequence according to the movement plan (Gálvez-García et al., 2014). The increased difficulty might inhibit the linking of bimanual sequences (Kennedy et al., 2021). As a result, movements might be preferentially represented discretely.

Second, linking the movements of two different body parts might be harder due to the organization and structure of the brain. Neural patterns pertaining to a bimanual movement sequence are spread out over both hemispheres due to the lateralisation of the motor cortex (Gerloff and Andres, 2002). Thus, a wider and bihemispheric network of brain modules is involved in bimanual compared to unimanual movement sequences, which might, accordingly, be more difficult to establish and maintain (Noble et al., 2014).

Third, neural crosstalk might interfere with learning of bimanual tasks (Kennedy et al., 2021; Swinnen, 2002). Neural crosstalk occurs during bimanual movements when neural signals designated to muscles in one arm are also sent to homologous muscles in the other arm (Cardoso de Oliveira, 2002). Interference emerges when additional, conflicting signals are received in close temporal proximity. In our active bimanual sequence group, left and right hand reaches had to be made directly following one another; thus the related neural signals may have resulted in interference, hampering adaptation.

### Sensory information as a substitute for active movement in sequence learning

In our study, providing prior visual or proprioceptive feedback of the opposite-arm movement did not enhance adaptation of the moving arm. This suggests that sensory information on its own does not provide a substitute for active movement in bimanual sequence learning. This finding contrasts with prior research in which visual and passive prior same-arm movements were effective cues for adaptation to opposing force fields (Howard et al., 2012). The contrast could be explained by sensory information of the same limb being weighted differently than information of another limb during sensorimotor integration processes. Sensorimotor integration is the ability to extract relevant sensory inputs to create informed motor outputs (Wolpert et al., 1995). Sensory information received from a same-arm visual or passive ”movement” directly affects the state estimation of the arm and thus the internal model and motor command of the next reach, allowing force field specific adaption. Sensory changes in another limb, however, do not bear the same relevance for the execution of a reach and may not be integrated computationally within a bimanual motor sequence. In addition, in the visual bimanual sequence group, participants may not have represented the red cursor as their left arm. In line with this explanation, visual cues which cannot be directly related to the state of the moving arm have not lead to adaptation in previous research, for instance, spatially static visual cues or field-specific cursor/background colors (Cothros et al., 2009; Howard et al., 2012, 2013).

Even though active engagement seems to be necessary for strong linking to occur across arms, some participants in the visual and passive bimanual sequence group were able to reduce their movement error over the course of the adaptation phase, indicating that inter-individual differences exist in whether and how cues can be used for motor adaptation. One factor that could influence individual adaptation is the attention given to the perceptual information of the opposite-arm. However, in our debriefing, four participants indicated that they had primarily attended to the color switch of the middle targets, rather than anticipating the end of the visual or proprioceptive left hand movement. Despite this strategy, these participants’ performance was not appreciably different from that of other participants. Thus, which factors may drive the adaptation differences between participants remains unclear.

In addition, we asked whether participants could transfer any learning obtained in a setting where the prior movement was only visual or passive to an active bimanual movement. This question was motivated by sports and rehabilitation practices, where a similar transfer would be highly desirable to support motor learning processes. In our experiment, however, learning was, for the most part, absent when only visual or passive-proprioceptive information of the prior opposite-arm movement were available. A transfer of overt improved performance to an active bimanual sequence was therefore impossible.

Surprisingly, there was immediate improvement in performance in the passive bimanual sequence group once they actively performed the task. The reduction of the MPE in the re-adaptation phase could, however, originate not from force field specific adaptation but from increased muscle co-contraction in the right arm. The increased stiffness would result in smaller MPEs without true learning of the force fields. Co-contraction of the right arm might be more pronounced in the re-adaptation than the adaptation phase because participants had to control their left arm movement in addition to their right arm during the active re-adaptation. Taken together, we cannot draw strong conclusions about transfer ability from linked sensory/active movements to active motor sequences.

### Motor adaptation vs. sequence learning

Motor theories usually distinguish between motor adaptation and skill learning. Motor adaptation entails a trial-by-trial change evoked by a mismatch between expected and received feedback and is thus a recalibration process (Wolpert et al., 2011). In contrast, skill learning entails the creation of a new movement pattern (Diedrichsen and Kornysheva, 2015). It is currently unknown whether findings about motor adaptation generalize to motor skill learning. In our view, it is crucial for complex bilateral motor behavior and learning to adjust part of a movement sequence according to kinematic parameters of the same sequence; this appears to us to be equivalent in multi-limb adaptation and skill learning. Our present results suggest, for example, that a novice juggler will improve over time by utilizing kinematic information from one arm to adjust movements of the other more and more. Bimanual tasks with concurrent movements of the arms have already shown that internal models of arm movements encompass not only kinematic information about the relevant arm but also about the opposite-arm to allow smooth compensation and flexible interaction between them (Yokoi et al., 2011). In our study, we provide first evidence that the selection process of internal models is influenced by prior movements of another limb and thereby contribute to a mechanistic understanding of complex bilateral motor behavior. Our finding is relevant for motor learning in both rehabilitation and sport settings. For example, in rehabilitation of stroke, activating a specific motor memory of the affected hand (e.g., reaching for a cup) could be cued and facilitated by a prior opposite-arm movement. Similarly, in sports, deliberately using movement sequences to differentiate between otherwise interfering moves (e.g., twisting once or twice in gymnastics) could be especially beneficial. These exciting prospects await future research.

## Acknowledgements

The authors would like to thank Pei-Cheng Shih for valuable discussion and help with programming and setting up the experiment, Philipp Kaniuth for useful comments on an earlier version of the manuscript, and Lisa Franke for help with data collection.

